# An integrative pan-cancer investigation reveals common genetic and transcriptional alterations of AMPK pathway genes as important predictors of clinical outcomes across major cancer types

**DOI:** 10.1101/735647

**Authors:** Wai Hoong Chang, Alvina G. Lai

## Abstract

The AMP-activated protein kinase (AMPK) is an evolutionarily conserved regulator of cellular energy homeostasis. As a nexus for transducing metabolic signals, AMPK cooperates with other energy-sensing pathways to modulate cellular responses to metabolic stressors. With metabolic reprogramming being a hallmark of cancer, the utility of agents targeting AMPK has received continued scrutiny and results have demonstrated conflicting effects of AMPK activation in tumorigenesis. Harnessing multi-omics datasets from human tumors, we seek to evaluate the seemingly pleiotropic, tissue-specific dependencies of AMPK signaling dysregulation. We interrogated copy number variation and differential transcript expression of 92 AMPK pathway genes across 21 diverse cancers involving over 18,000 patients. Cox proportional hazards regression and receiver operating characteristic analyses were used to evaluate the prognostic significance of AMPK dysregulation on patient outcomes. A total of 24 and seven AMPK pathway genes were identified as having loss- or gain-of-function features. These genes exhibited tissue-type dependencies, where survival outcomes in glioma patients were most influenced by AMPK inactivation. Cox regression and log-rank tests revealed that the 24-AMPK-gene set could successfully stratify patients into high- and low-risk groups in glioma, sarcoma, breast and stomach cancers. The 24-AMPK-gene set could not only discriminate tumor from non-tumor samples, as confirmed by multidimensional scaling analyses, but is also independent of tumor, node and metastasis staging. AMPK inactivation is accompanied by the activation of multiple oncogenic pathways associated with cell adhesion, calcium signaling and extracellular matrix organization. Anomalous AMPK signaling converged on similar groups of transcriptional targets where a common set of transcription factors were identified to regulate these targets. We also demonstrated crosstalk between pro-catabolic AMPK signaling and two pro-anabolic pathways, mammalian target of rapamycin and peroxisome proliferator-activated receptors, where they act synergistically to influence tumor progression significantly. Genetic and transcriptional aberrations in AMPK signaling have tissue-dependent pro- or anti-tumor impacts. Pan-cancer investigations on molecular changes of this pathway could uncover novel therapeutic targets and support risk stratification of patients in prospective trials.

## Introduction

The AMP-activated protein kinase (AMPK) is an evolutionary conserved key player responsible for energy sensing and homeostasis. Orthologous copies of AMPK prevail universally as heterotrimeric complexes where the human genome encodes two genes for the α catalytic subunit, two β regulatory subunit genes and three γ subunit genes. Historically, AMPK was discovered as a crucial regulator of lipid metabolism(1). Since then, AMPK is implicated in a wide variety of fundamental metabolic processes as well as in metabolic diseases such as cancer and diabetes(2). The first link between AMPK and cancer was identified through the tumor-suppressive function of LKB1, which is upstream of the mTOR pathway(3). The tumor-suppressive roles of AMPK were pharmacologically demonstrated by the application of metabolic inhibitors such as the antidiabetic metformin and the mimetic of AMP, AICAR(4–6). Numerous studies have since compellingly established the promiscuous nature of these pharmacological agents, whereby the inhibition of cancer cell proliferation occurs through non-specific AMPK-independent avenues(7, 8).

In contrast to the tumor-suppressive results from pharmacological studies, genetic experiments on cancer cells have credibly demonstrated that AMPK activation is crucial for tumor progression and survival(9–12). A myriad of metabolic stressors, such as oxygen deprivation, nutrient starvation and oxidative stress, exists within the tumor microenvironment. Metabolic reprogramming during carcinogenesis would thus trigger AMPK activation to enable cells to survive under conditions of stress typically found in the tumor microenvironment, hence conferring an overall tumor-promoting effect. AMPK is also shown to support cancer growth and migration through crosstalk with other pro-oncogenic pathways. For instance, overexpression of oncogenes MYC and SRC or the loss of the tumor suppressor folliculin could lead to AMPK activation(13–17).

Genetic and pharmacological studies have paved the way for our understanding of the function of AMPK in cancer. However, anti- and pro-neoplastic features of AMPK remain controversial potentially due to the oversimplification of AMPK-modulated processes in in vitro and non-human in-vivo models. The genetic and clinical landscape of AMPK signaling has not been systematically investigated. Thus, our study aims to address an unmet need to rigorously investigate the role of AMPK in diverse cellular context using multi-omics data from actual tumors where we examined somatic copy number alterations, transcriptional and clinical profiles of tumors from 21 cancer types. Our analyses of clinical samples at scale would complement evidence from pharmacological and genetic studies to better elucidate the multi-faceted and cell-specific nature of AMPK signaling on tumor progression.

## Methods

### AMPK pathway genes and cancer cohorts

92 AMPK pathway genes were retrieved from the Kyoto Encyclopedia of Genes and Genomes (KEGG) database (Table S1). Clinical, genomic and transcriptomic datasets of 21 cancers involving over 18,484 patients were downloaded from The Cancer Genome Atlas (TCGA)(18).

### Copy number variation, differential expression, multidimensional scaling and survival analyses

Detailed methods of the above analyses were previously published and thus will not be repeated here as per the journal guidelines(19, 20). To summarize, discrete amplification and deletion indicators for copy number variation analyses were obtained from GISTIC gene-level tables(21). GISTIC values of +1 and −1 were annotated as shallow amplification and shallow deletion (heterozygous) events respectively. GISTIC values of +2 and −2 were annotated as deep amplification and deep (homozygous) deletion events respectively. Multidimensional scaling analyses and permutational multivariate analysis of variance (PERMANOVA) were performed using the R vegan package. Survival analyses were performed using Cox proportional hazards regression and the log-rank test. Sensitivity and specificity of the 24-AMPK-gene set were assessed using receiver operating characteristic analyses. Differential expression analyses were performed on patients stratified into high- (4th quartile) and low- (1st quartile) expressing groups using the 24-gene-set to determine the transcriptional effects of anomalous AMPK signaling.

### Pathway and transcription factor analyses

Genes that were differentially expressed (DEGs) between the 4th and 1st quartile patient groups were mapped to KEGG, Gene Ontology and Reactome databases using g:profiler(22) to ascertain biological processes and signaling pathways that were enriched. The Enrichr tool(23, 24) was used to map DEGs to the ChEA and ENCODE transcription factor (TF) databases to identify TFs that were significantly enriched as regulators of the DEGs.

### Calculating the 24-AMPK-gene score, peroxisome proliferator-activated receptors (PPAR) score and mammalian target of rapamycin (mTOR) score

AMPK scores were calculated from the mean expression of the following genes: SLC2A4, FOXO3, PPP2CB, PIK3CD, CAB39L, CCNA1, FBP1, FBP2, FOXO1, HMGCR, IRS2, PIK3R1, SIRT1, TBC1D1, PPARGC1A, PPP2R2C, MLYCD, PFKFB3, PPP2R2B, PRKAA2, LEPR, CAB39, IRS1 and PFKFB1. PPAR scores for each patient were calculated by taking the mean expression of PPAR signature genes: PLIN5, PPARG, ACADM, GK, CPT2, SCP2, ACAA1, APOA1, PPARA, ACOX2, ANGPTL4, FABP3, PLIN2, AQP7, ACSL1, FABP5, ACADL, and PCK2(20). mTOR/PI3K/AKT scores for each patient were calculated using the following equation: mTOR/PI3K/AKT score = AKT + mTOR + GSK3 + S6K + S6 – PTEN(25).

## Results

### Pan-cancer genomic and transcriptional alterations of AMPK pathway genes

Focusing on the genomic and transcriptomic landscape of 92 genes associated with AMPK signaling retrieved from KEGG across 21 cancer types involving 18,484 patients (Table S1), we interrogated somatic copy number alterations (SCNA) and mRNA expression. To determine the effects of genomic alterations in AMPK pathway genes, we classified genes as having high-level amplifications (gains), low-level amplifications, deep (homozygous) deletions and shallow (heterozygous) deletions. To evaluate pan-cancer patterns of SCNAs, we considered genes that were gained or lost in at least 20% of samples within a cancer type and in at least one-third of cancer types, i.e., at least seven cancer types. A total of 46 genes were recurrently amplified, while 49 genes were recurrently lost (Figure 1; Table S2). AMPK is the central regulator of cellular energy levels, which controls a number of downstream targets, an example being the nuclear receptor HNF4A. Remarkably, HNF4A was found to be the most amplified gene; identified as being recurrently amplified in >20% of samples in all 21 cancers (Figure 1; Table S2). This is followed by CFTR (18 cancer types) and four other genes that were amplified in 17 cancer types (ADIPOR2, LEP, PRKAG2 and RHEB) (Figure 1; Table S2). In contrast, PPP2R2A was the most deleted gene found in >20% of samples across 17 cancers, followed by the deletion of SLC2A4 in 16 cancers and five additional genes (FOXO3, PPP2CB, PPP2R2D, PPP2R5C and PPP2R5E) in 15 cancer types (Figure 1; Table S2). Among all cancer types, the highest number of amplified AMPK pathway genes was observed in esophageal carcinoma (ESCA; 44 genes) followed by bladder cancer (BLCA; 42 genes) and lung cancer (41 genes in both lung squamous cell carcinoma [LUSC] and adenocarcinoma [LUAD]) (Figure 1). Glioma tumors (GBMLGG), in contrast, had only five genes that were recurrently amplified (Figure 1). In terms of somatic deletions, LUSC and ESCA both had 49 genes deleted while no recurrent deletions were observed in papillary renal cell carcinoma (KIRP) (Figure 1).

**Fig. 1.**
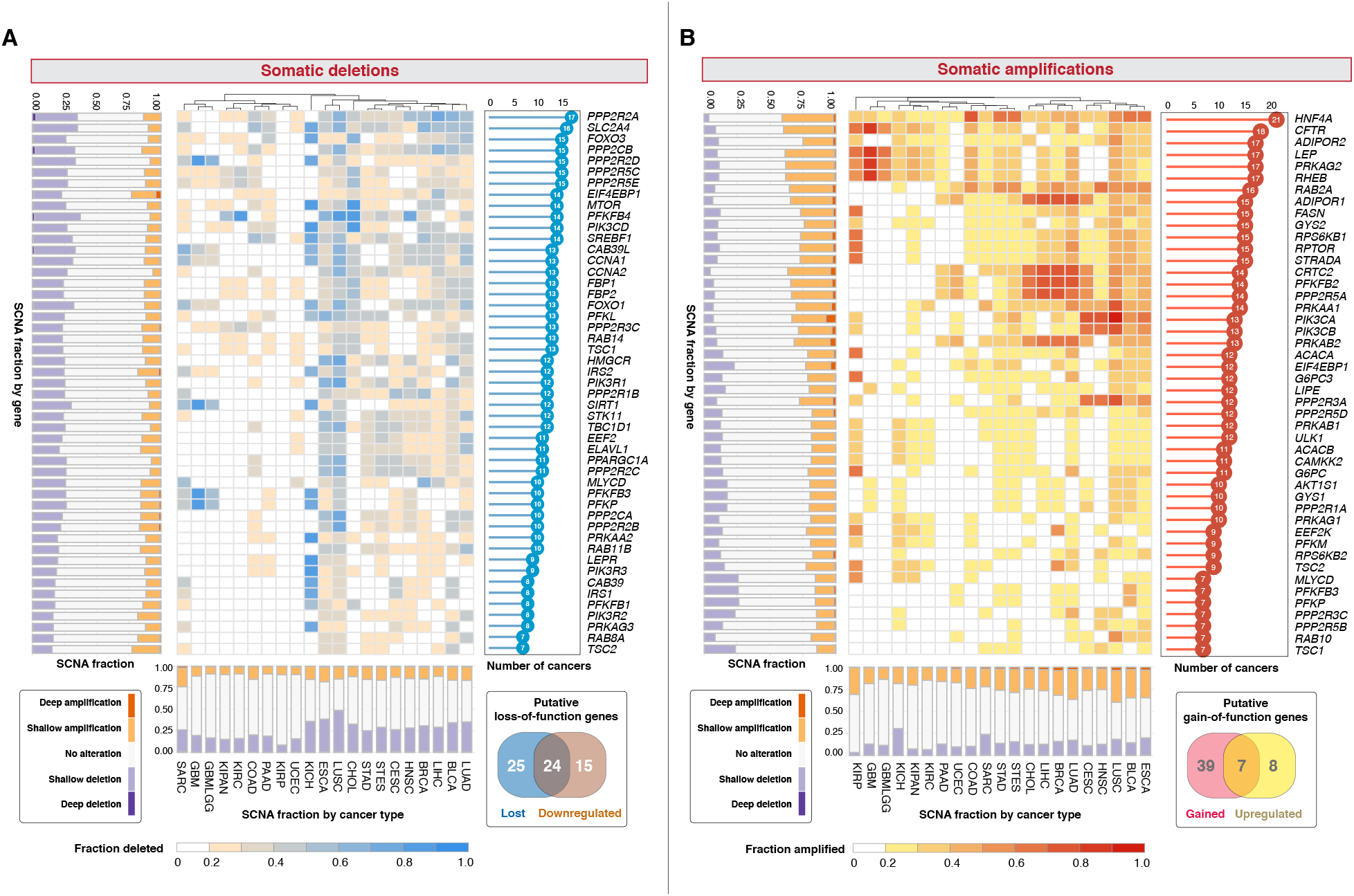
The landscape of somatic copy number alterations of AMPK pathway genes. Heatmaps depict (A) fraction of samples within each cancer type that harbor somatic deletions and (B) somatic amplifications. Forty-nine genes are recurrently deleted in at least 20% of tumors within each cancer and in at least seven cancer types. Forty-six genes are recurrently amplified in at least 20% of tumors within each cancer and in at least seven cancer types. Stacked bar charts on the y-axes illustrate the fraction of samples that possess copy number variation of a gene under consideration grouped by shallow and deep deletions or amplifications. Stacked bar charts on the x-axes illustrate the fraction of samples within each cancer type that contain shallow and deep deletions or amplifications. The bar charts on the right of each heatmap depict the number of cancer types with at least 20% of samples affected by gene deletions and amplifications. The Venn diagrams demonstrate the identification of 24 putative loss and seven gain-of-function genes from gene sets that are somatically altered and differentially expressed. Cancer cohorts analyzed with corresponding TCGA abbreviations are listed in parentheses: bladder urothelial carcinoma (BLCA), breast invasive carcinoma (BRCA), cervical squamous cell carcinoma and endocervical adenocarcinoma (CESC), cholangiocarcinoma (CHOL), colon adenocarcinoma (COAD), esophageal carcinoma (ESCA), glioblastoma multiforme (GBM), glioma (GBMLGG), head and neck squamous cell carcinoma (HNSC), kidney chromophobe (KICH), pan-kidney cohort (KIPAN), kidney renal clear cell carcinoma (KIRC), kidney renal papillary cell carcinoma (KIRP), liver hepatocellular carcinoma (LIHC), lung adenocarcinoma (LUAD), lung squamous cell carcinoma (LUSC), pancreatic adenocarcinoma (PAAD), sarcoma (SARC), stomach adenocarcinoma (STAD), stomach and esophageal carcinoma (STES) and uterine corpus endometrial carcinoma (UCEC).

We reasoned that SCNAs associated with transcriptional alterations could be considered as putative gain- or loss-of-function events. Differential expression analyses between tumor and non-tumor samples in each cancer revealed that 15 and 39 genes were significantly upregulated and downregulated in at least seven cancer types respectively (Figure S1). Of these differentially expressed genes, seven and 24 genes were also recurrently amplified and deleted respectively (Venn diagram in Figure 1). Both gene sets were mutually exclusive, i.e., the genes either had gain-or-function or loss-of-function features, but not both.

### Molecular underpinnings of patient survival involving putative loss-of-function AMPK pathway components

We next investigated the impact of transcriptional dysregulation of the putative gain- and loss-of-function genes identified previously on patient survival outcomes across all cancer types. Employing Cox proportional hazards regression, we observed that all 31 genes (seven gain-of-function and 24 loss-of-function genes), were prognostic in at least one cancer type (Figure 2A). The highest number of prognostic genes was observed in glioma (GBMLGG) tumors (26/31 genes), while none of the 31 genes were significantly associated with overall survival outcomes in ESCA and cholangiocarcinoma (CHOL) (Figure 2A). Intriguingly, although ESCA had the highest number of SCNAs (Figure 1), none of the genes harbored prognostic information, suggesting that alterations in AMPK signaling components have minimal roles in driving tumor progression and patient outcomes. FBP1 was significantly associated with overall survival outcomes in 10 cancers while PPP2R2C and PPP2R2B in 8 cancers (Figure 2A). FBP2 is the least prognostic gene in only one cancer type, cervical squamous cell carcinoma and endocervical adenocarcinoma; CESC (Figure 2A).

**Fig. 2.**
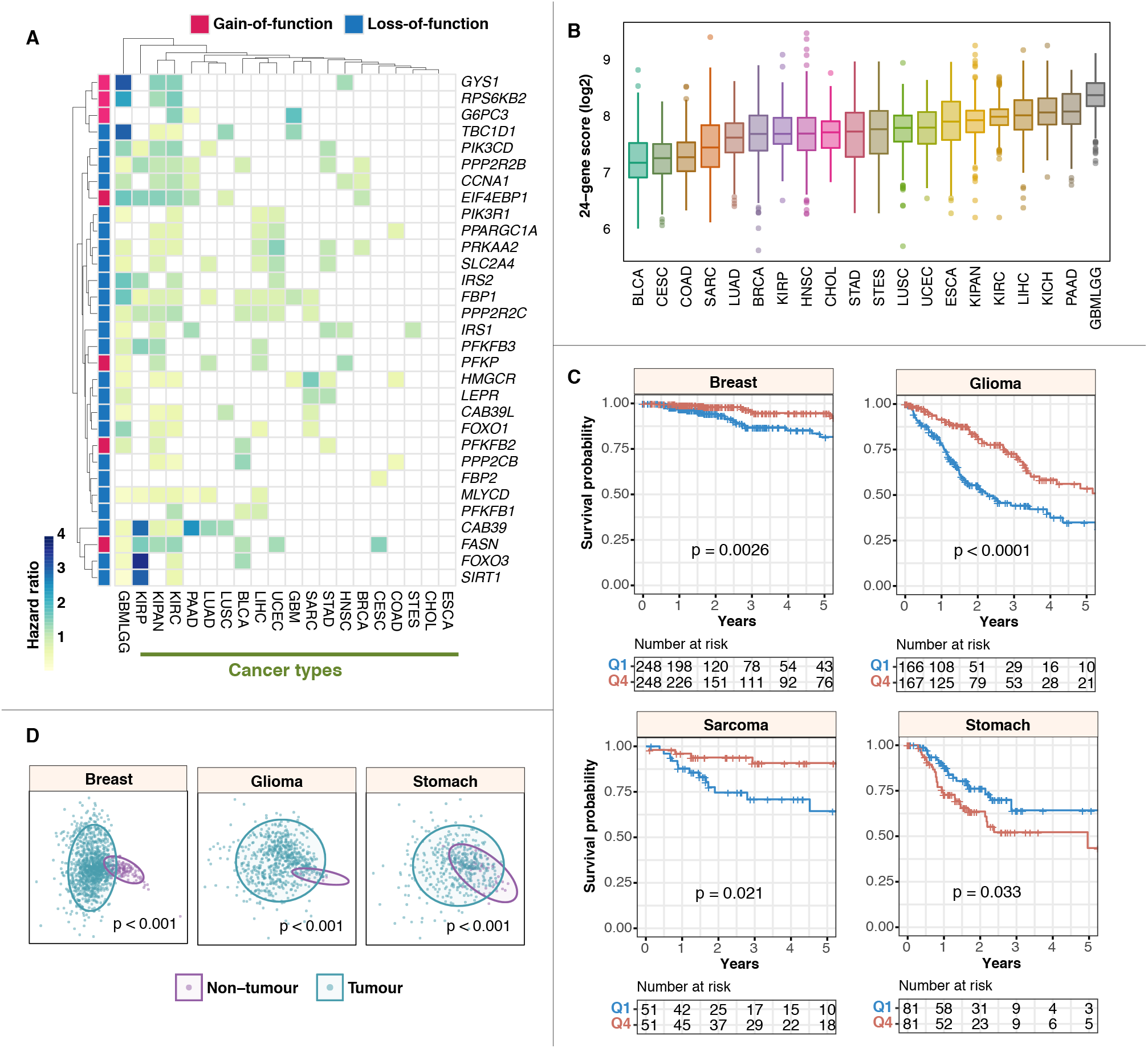
Prognostic significance of AMPK loss- and gain-of-function genes. (A) Heatmap illustrates significant hazard ratio values from Cox proportional hazards regression analyses on the 24 loss-of-function and seven gain-of-function genes across all cancers. (B) The distributions of 24-AMPK-gene scores in each cancer are illustrated in the boxplot. Cancers are sorted from low to high median scores. Refer to Figure 1 legend for cancer abbreviations. (C) Kaplan-Meier analyses and log-rank tests revealed the prognostic significance of the 24-AMPK-gene set in four cancer types. Patients are stratified into Q1 (1st quartile) and Q4 (4th quartile) groups based on their 24-gene scores for log-rank tests. (D) Multidimensional scaling analyses of the 24-gene set depicted in 2-dimensional space. Significance differences in the distribution between tumor and non-tumor samples are confirmed by PERMANOVA.

Given the prevalence of loss-of-function phenotypes in determining clinical outcomes (Figure 2A), we proceeded to examine the combined impact of all 24 loss-of-function genes on patient survival and oncogenic dysregulation. To determine the extent of AMPK pathway variation across the 21 cancers, we calculated ‘pathway scores’ for each of the 18,484 tumor samples by taking the mean transcript expression values of the 24 genes: SLC2A4, FOXO3, PPP2CB, PIK3CD, CAB39L, CCNA1, FBP1, FBP2, FOXO1, HMGCR, IRS2, PIK3R1, SIRT1, TBC1D1, PPARGC1A, PPP2R2C, MLYCD, PFKFB3, PPP2R2B, PRKAA2, LEPR, CAB39, IRS1 and PFKFB1. We observed interesting patterns when cancers were ranked from low to high, based on their median pathway scores (Figure 2B). GBMLGG had the highest median pathway score, while BLCA and CESC were found at the lower end of the spectrum (Figure 2B). As expected, Kaplan-Meier analysis revealed a significant difference in overall survival between glioma patients (P < 0.0001) stratified by low and high 24-gene pathway scores (Figure 2C). Interestingly, the contribution of AMPK signaling in cancer prognostication is cancer-type dependent. As in glioma, log-rank tests revealed that patients with high 24-gene scores had significantly improved survival outcomes in breast cancer (P = 0,0026) and sarcoma (P = 0.021) (Figure 2C). In contrast, high expression of the 24 genes was associated with increased mortality rates in stomach adenocarcinoma (P=0.033) (Figure 2C). These results were in agreement when independently validated using the Cox regression approach: breast (hazard ratio [HR] = 0.397; P = 0.0028), glioma (HR = 0.430; P < 0.0001), sarcoma (HR = 0.379; P = 0.021) and stomach (HR = 1.825; P = 0.034) cancers (Table S3). Since the 24-gene score could be used to stratify patients into high- and low-risk groups, we predict that when considered together, gene expression values could discriminate tumor from non-tumor samples. Although analysis could not be performed on sarcoma (this dataset only had two non-tumor samples), multidimensional scaling analyses and PERMANOVA tests of breast (P < 0.001), glioma (P < 0.001) and stomach (P < 0.001) cancers revealed significant separation between tumor and non-tumor samples in two-dimensional space (Figure 2D). Overall, this suggests that the 24-gene set could be harnessed as a diagnostic biomarker for early cancer detection.

To determine the independence of the 24-gene set from other clinicopathological features, we employed multivariate Cox regression and observed that the 24-gene set is independent of tumor, node and metastasis (TNM) staging (where available) in breast (HR = 0.403; P = 0.0043) and stomach cancers (HR = 1.835; P = 0.038) (Table S3). Similarly, Kaplan-Meier analyses and log-rank tests confirmed that the 24-gene set allowed further risk stratification of patients with tumors of the same TNM stage: breast (P < 0.0001) and stomach (P = 0.022) (Figure 3A). Furthermore, we observed that within a histological subtype of sarcoma, leiomyosarcoma, patients with elevated AMPK signaling had significantly better survival outcomes (P = 0.0072) (Figure 3A); consistent with our previous observation that high pathway scores were associated with good prognosis in sarcoma (Figure 2C).

**Fig. 3.**
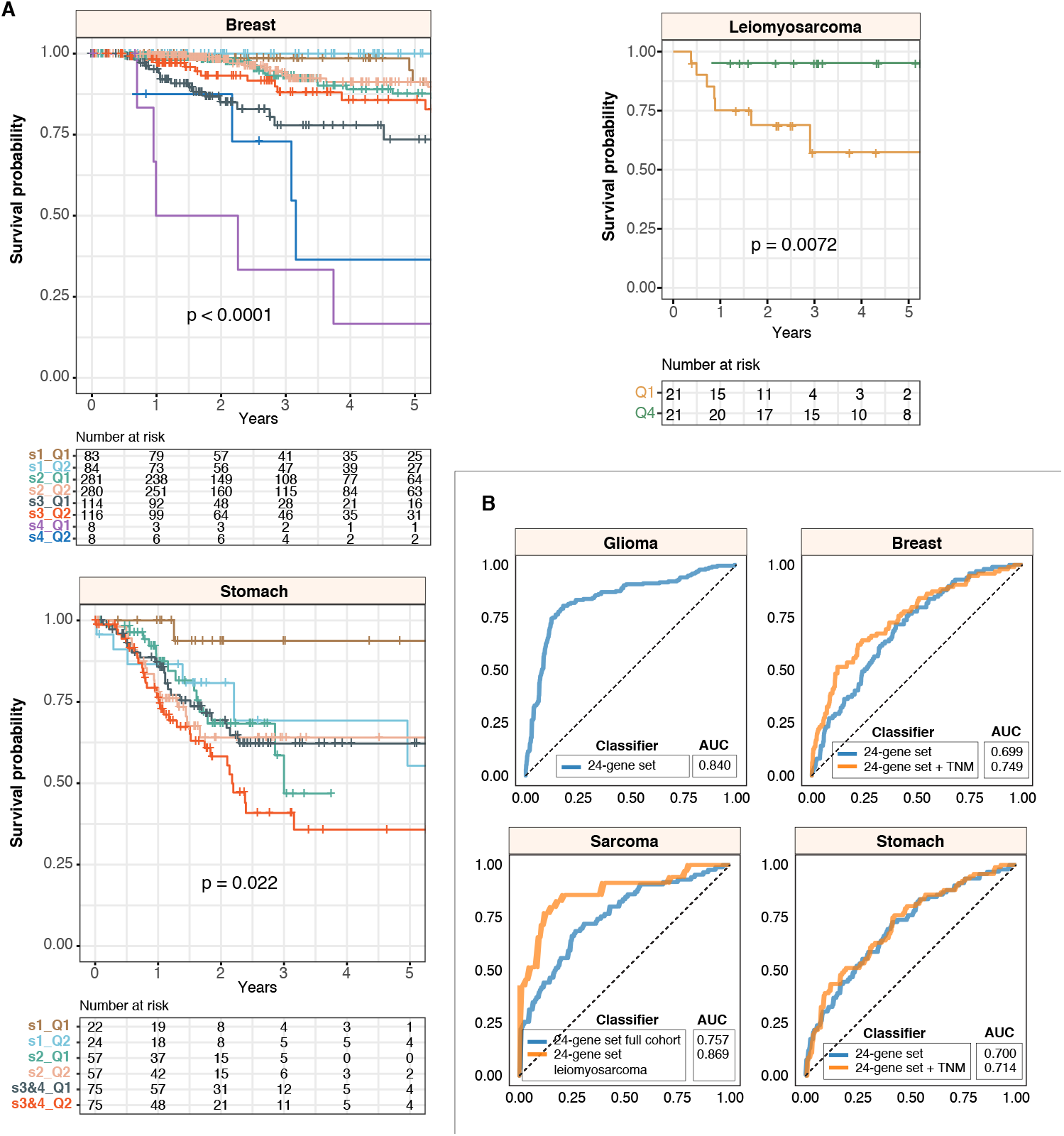
The 24-AMPK-gene set is independent of tumor stage and histological subtype. (A) Kaplan-Meier analyses of patients grouped by tumor, node and metastasis (TNM) stage (breast and stomach cancers) or by the histological subtype of leiomyosarcoma and the 24-gene score. For leiomyosarcoma, the log-rank test reveals a significant difference in survival rates between 1st and 4th quartile patients. (B) Receiver-operating characteristic (ROC) analyses on the 5-year predictive performance of the 24-gene set. ROC curves generated by the 24-gene set are compared to curves generated from both 24-gene set and TNM staging, where available, or histological subtype. AUC: area under the curve.

We next explored the predictive performance (sensitivity versus specificity) of the 24-gene set in all four cancer types using receiver operating characteristic analysis. The area under the ROC curve (AUC) is an indication of how well the gene set could predict patient survival, which ranges from 0.5 to 1. We found that the combined model uniting both 24-gene set and TNM staging outperformed the 24-gene set when considered on its own in breast cancer patients (AUC = 0.749 vs. 0.699) (Figure 3B). Since the AUC for TNM staging was only marginally higher than 0.5 in stomach cancer (AUC = 0.561) while the AUC for the 24-gene set was 0.700, TNM staging did not contribute to any increase in performance of the gene set (Figure 3B). AUCs of the 24-gene set in glioma and sarcoma were 0.840 and 0.757 respectively (Figure 3B). Within the leiomyosarcoma histological subtype, AUC was even higher at 0.869 (Figure 3B).

### Oncogenic transcriptional alterations associated with AMPK pathway inactivation

AMPK pathway inactivation was associated with altered survival outcomes in patients (Figure 2 and 3). We predict that this could be due to broad transcriptional dysregulation arising from abnormal AMPK signaling. To investigate this phenomenon, we performed differential expression analyses between patients stratified by the 24-gene set into high (4th quartile) and low (1st quartile) expression groups and found that an outstanding number of 122 common genes that were significantly differentially expressed in all four cancer types (Figure 4A). The highest number of differentially expressed genes (DEGs) was observed in stomach cancer (2,496 genes), followed by sarcoma (1,842 genes), glioma (1,523 genes) and breast cancer (1,086 genes) (Figure 4A; Table S4). The DEGs were mapped to KEGG, Gene Ontology and Reactome databases to determine whether they were associated with any functionally enriched pathways. Intriguingly, all four cancer types share similar patterns of functional enrichments (Figure 4B and 4C). For instance, biological processes associated with cell communication, signal transduction, cell differentiation, cell signaling, cell adhesion and cell morphogenesis were enriched in all four cancers (Figure 4C). In terms of specific signaling pathways, calcium signaling, cAMP signaling, and processes associated with extracellular matrix organization were among the most enriched (Figure 4C).

**Fig. 4.**
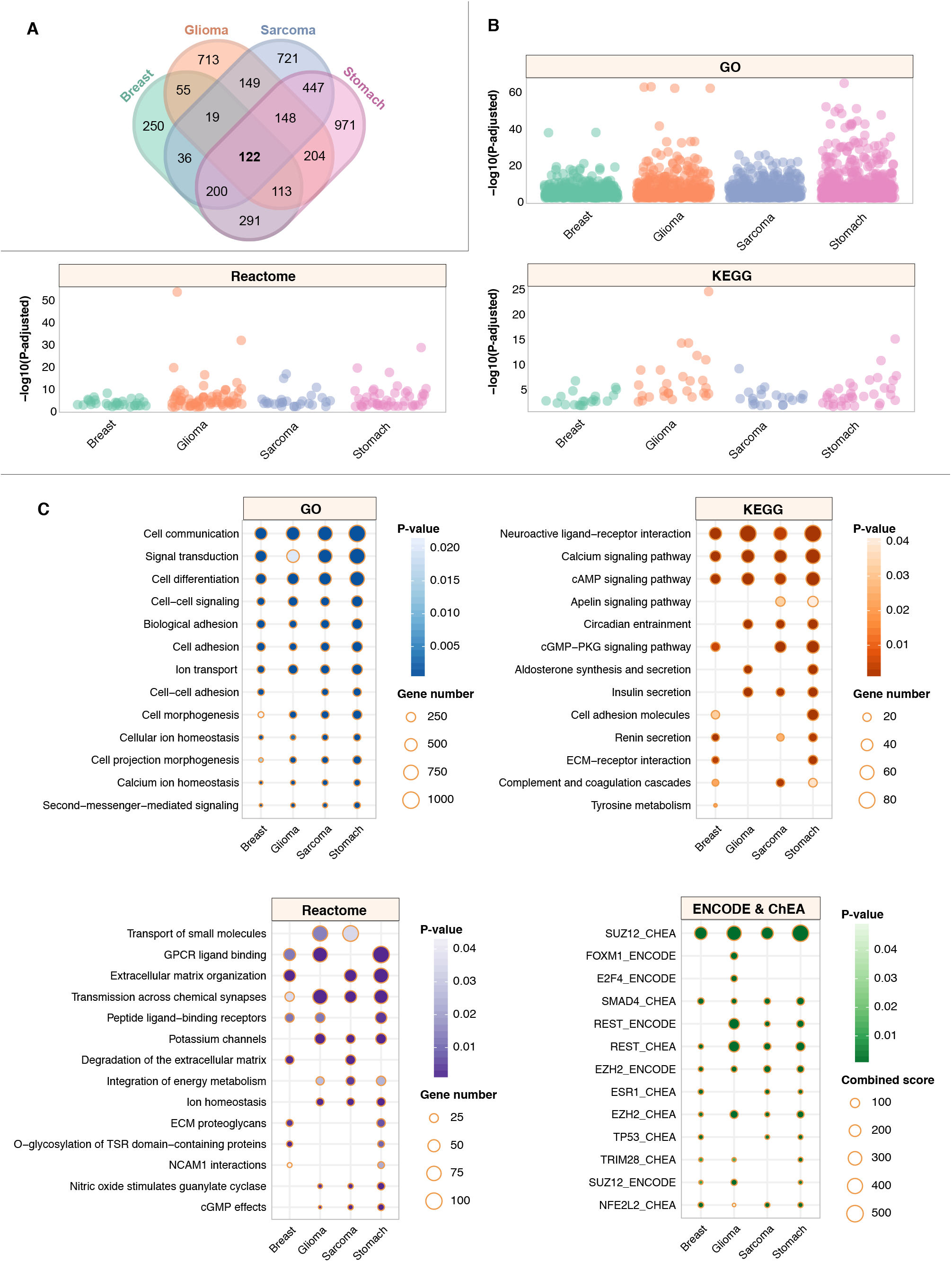
AMPK inactivation drives oncogenic transcriptional alterations in diverse biological processes and signaling modules. (A) Venn diagram illustrates the number of differentially expressed genes (DEGs) between 1st and 4th quartile patients, as stratified using the 24-AMPK-gene set, in four cancer types. A total of 122 DEGs were common in all four cancers. (B) Dot plots depict the number of significantly enriched pathways and biological processes upon the mapping of DEGs to KEGG, Gene Ontology and Reactome databases. Each dot represents an enriched event. (C) Ontologies that exhibit similar patterns of enrichment across four cancers are shown. DEGs are also mapped to ENCODE and ChEA transcription factor (TF) databases to determine enriched TF binding associated with DEGs.

To further identify potential transcriptional regulators of the DEGs, we mapped the DEGs to ENCODE and ChEA transcription factor (TF) binding databases. Remarkably, we identified common TFs, shared across all four cancers, that were implicated as direct binding partners of the DEGs (Figure 4C). Five TFs, SUZ12, SMAD4, REST, EZH2 and NFE2L2, were found to be enriched in all four cancers, suggesting that transcriptional dysregulation of tumors with aberrant AMPK signaling involved direct physical associations of these TFs with target DEGs (Figure 4C). Curiously, FOXM1 and E2F4 were enriched only in glioma tumors, which deserves further exploration in the next section. Overall, our analyses demonstrated that impaired AMPK signaling resulted in common patterns of oncogenesis, which affect the severity of cancer and consequently, mortality rates in patients.

### Downstream targets of EZH2, NFE2L2, REST, SMAD4 and SUZ12 were associated with survival outcomes

Pathways modulating energy homeostatic may transduce signals to influence other cognate signaling modules. EZH2, NFE2L2, REST, SMAD4 and SUZ12 were all implicated as common transcriptional regulators of DEGs in glioma, sarcoma, breast and stomach cancers, suggesting that altered AMPK signaling converged on similar groups of transcriptional targets. Of all the target DEGs of the aforementioned TFs, 8, 10, 24, 12 and 48 genes were found to be common targets of EZH2, NFE2L2, REST, SMAD4 and SUZ12 respectively in all four cancers (Figure 5A). Concatenating all five gene sets yielded 71 unique genes, i.e., genes that were binding targets of more than one TF were considered only once. To gain further insights into how AMPK inactivation influences tumor progression, we performed Cox regression analyses to determine the association between each of the 71 genes and survival outcomes. The highest number of prognostic genes was observed in glioma; 66 genes (61 good prognoses and five adverse prognoses) (Figure 5B). In contrast, 54 out of 71 genes were associated with adverse prognosis in stomach cancer (Figure 5B). These observations were consistent with the 24-AMPK-gene set being positive and negative prognostic factors in glioma and stomach cancer respectively (Figure 2), which mirrored the behavior of DEGs identified as a result of aberrant AMPK signaling (Figure 4C). Of the 71 genes, only 15 and ten were significantly associated with survival outcomes in sarcoma and breast cancer respectively (Figure 5B). Collectively, our results suggest that the AMPK pathway and its interaction with other signaling modules are key determinants of patient outcomes in multiple cancer types.

**Fig. 5.**
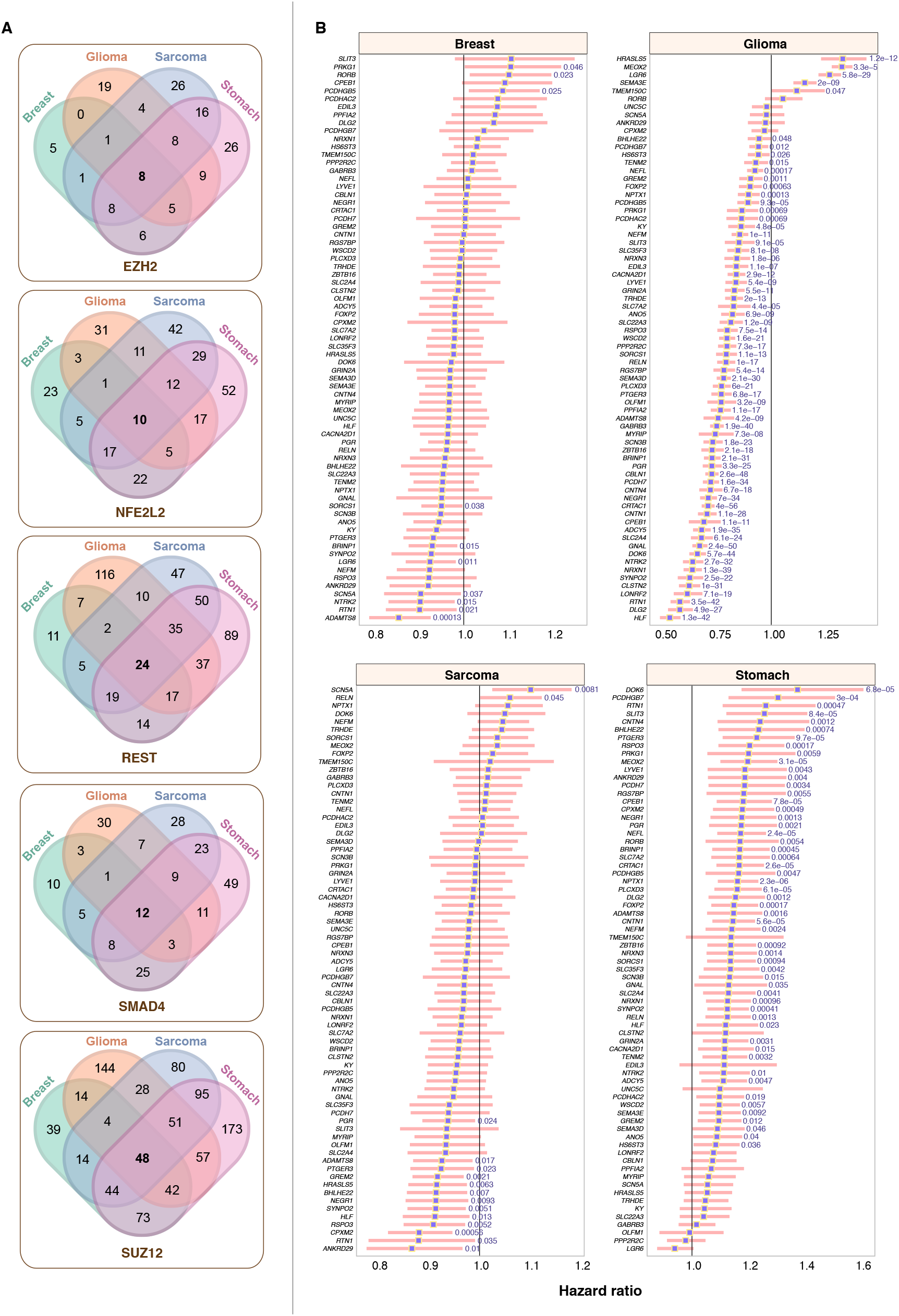
Prognostic significance of DEGs targeted by enriched TFs. (A) Venn diagrams illustrate the extent of overlap between DEGs targeted by EZH2, NFE2L2, REST, SMAD4 and SUZ12 across four cancers. (B) Forest plots depict DEGs that are significantly associated with overall survival outcomes. Hazard ratios are denoted as purple squares while pink bars represent the 95% confidence intervals. Significant Wald test P values are indicated in blue.

### Prognostic significance of joint AMPK pathway activity and transcriptional levels of five oncogenic TFs in patients with glioma

Having discovered the importance of the 24-AMPK gene set, we sought to explore the crosstalk between AMPK signaling and TF activity in glioma. As previously mentioned, glioma had the highest 24-AMPK-gene score (Figure 2B) with a vast majority of the genes conferring prognostic information (Figure 2A). Moreover, 66 of the 71 transcriptional targets of the five common TFs identified in patients with altered AMPK signaling were significantly associated with survival outcomes in glioma (Figure 5B). Additionally, TFs FOXM1 and E2F4 were identified to be enriched only in glioma tumors (Figure 4C). Thus, we predict that a joint model uniting AMPK and TF expression profiles would allow further delineation of patients into additional risk groups and if so, allowing combined targeting of AMPK and candidate TFs for therapeutic action. As done previously, we calculated AMPK scores for each patient based on the mean expression of the 24 genes. Interestingly, we found that AMPK scores were significantly negatively correlated with TF expression levels in glioma: E2F4 (rho = −0.48, P 0.0001), EZH2 (rho = −0.57, P < 0.0001), FOXM1 (rho = −0.49, P < 0.0001), SMAD4 (rho = −0.18, P < 0.0001) and SUZ12 (rho = −0.23, P < 0.0001) (Figure 6A). We subsequently categorized patients into four groups using the median cutoff of the AMPK scores and TF expression values: 1) low-low, 2) high-high, 3) low AMPK score and high TF expression and 4) high AMPK score and low TF expression. Log-rank tests revealed that patients stratified into the four groups had survival rates that were significantly different: E2F4 (P < 0.0001), EZH2 (P < 0.0001), FOXM1 (P < 0.0001), SMAD4 (P < 0.0001) and SUZ12 (P < 0.0001) (Figure 6B). For E2F4, EZH2, FOXM1 and SUZ12, patients with low AMPK scores and high TF expression performed the worst: E2F4 (HR = 3.916; P < 0.0001), EZH2 (HR = 4.004; P < 0.0001), FOXM1 (HR = 5.268; P < 0.0001) and SUZ12 (HR = 2.197; P < 0.0001) (Figure 6C). For SMAD4, patients within the low-low category had the highest mortality rates (HR = 3.326; P < 0.0001) (Figure 6C).

**Fig. 6.**
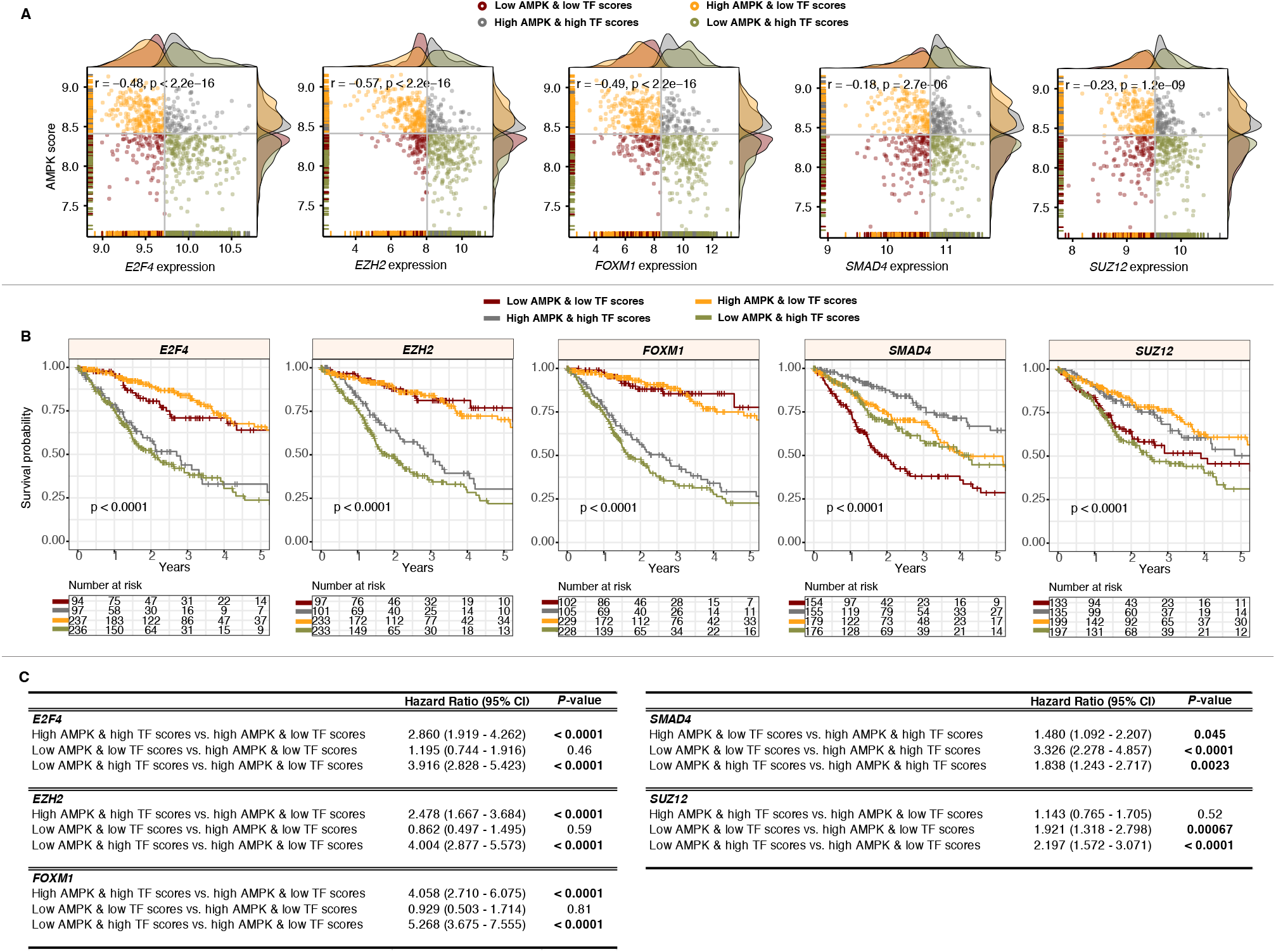
Prognostic relevance of candidate TFs and the 24-AMPK-gene set in glioma. (A) Scatter plots illustrate significant negative correlations between AMPK scores and TF expression levels in glioma. Patients are separated and color-coded into four categories based on median AMPK and TF scores. Density plots appended to the y- and x-axes demonstrate the distribution of AMPK and TF scores. (B) Log-rank tests are performed on the four patient groups to demonstrate the utility of combined AMPK and TF scores in patient stratification. (C) Univariate Cox regression analyses are performed to compare patient groups where significant P values are highlighted in bold. CI: confidence interval.

### Crosstalk between AMPK and other anabolic-related pathways, PPAR and mTOR

AMPK’s anti-anabolic and pro-catabolic activities may work in concert with other metabolic pathways. To investigate the synergistic effects of AMPK and two pro-anabolic pathways, peroxisome proliferator-activated receptors (PPAR) and mammalian target of rapamycin (mTOR) signaling in tumor progression, we calculated PPAR and mTOR pathway scores (detailed in the methods section) for each glioma tumor. Low AMPK scores were associated with poor outcomes in glioma (Figure 2). To evaluate AMPK and PPAR or mTOR as combined models, patients were separated into four groups using the median cutoff, as mentioned previously. Interestingly, when AMPK and PPAR scores were collectively used for patient stratification, we found that patients with low AMPK and high PPAR scores had the highest death rates (HR = 11.308, P < 0.0001), confirming that PPAR hyperactivation is associated with poor outcomes in glioma tumors with low AMPK activity(20) (Figure 7). In contrast, when considering mTOR activity, patients with low AMPK and low mTOR scores performed the worst (HR = 3.023, P < 0.0001) (Figure 7). The results overall suggest that the AMPK pathway could act synergistically with PPAR and mTOR signaling to influence cancer progression significantly.

**Fig. 7.**
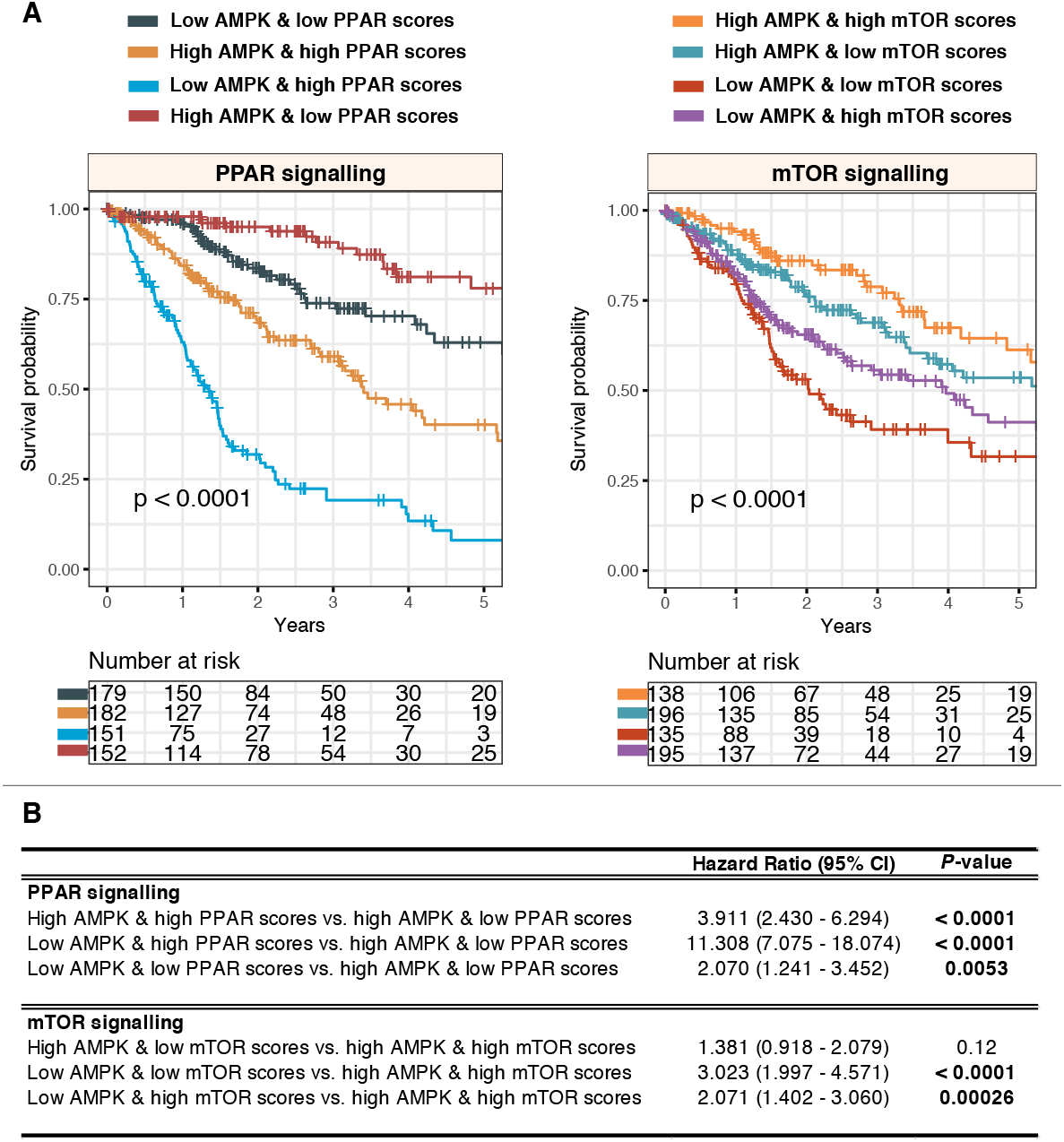
Crosstalk between AMPK signaling and PPAR or mTOR pathways in glioma. (A) Log-rank tests are performed on patient groups separated into four categories based on median AMPK and PPAR or mTOR scores. (B) Univariate Cox regression analyses are performed to compare patient groups where significant P values are highlighted in bold. CI: confidence interval.

## Discussion and Conclusion

While the role of AMPK in energy-sensing is well understood, its full potential in metabolic diseases such as cancer remains an open topic of debate. Despite extensive efforts spent on elucidating the role of AMPK signaling(2, 9, 11), there remains no consensus on whether AMPK promotes or suppresses tumor progression. Exploiting a rich reservoir of pan-cancer datasets afforded to us by TCGA, we performed a thorough examination of genomic and transcriptomic profiles of 92 AMPK pathway genes in diverse cancer types. Our current understanding of AMPK signaling is fueled by genetic studies in cell lines and animal models(2). Although useful in determining causal relationships, results from in vitro cell lines and animal models may have limited translational relevance as they do not accurately reflect human pathology(26). Animal models may offer additional mechanistic insights, but limitations in ethics and costs remain. Moreover, the complexity of human cancers is not accurately modeled in animals; less than 8% of results from animal models are translated to clinical trials(27). Despite analyses on tumor genetic datasets providing mostly correlative outcomes, they remain valuable in understanding disease-specific molecular pathology when interrogated at scale on large patient groups(28), and when results are considered in relation to those obtained from cell lines and animal models.

Employing pan-cancer population data, our study identified conserved and unique patterns of AMPK signaling across diverse cancer types. Analyses at two molecular levels (genetic and transcriptional) yielded a more comprehensive depiction of AMPK signaling, where we identified genes that were both somatically altered and differentially expressed. These putative loss- or gain-of-function genes are more likely to impact tumor progression as they are altered at both macromolecular levels. As reported in other studies, we confirmed that AMPK signaling could either be oncogenic or tumor suppressive depending on the cellular context. Intuitively, since AMPK is anti-anabolic, its function may not be fitting for tumor growth and proliferation. This is consistent with reports demonstrating AMPK’s tumor suppressive activity(29, 30). A study on lymphoma demonstrates that AMPK downregulation induces the Warburg effect and hypoxia signaling in mice(31). AMPK is proposed to act as a metabolic gatekeeper to limit cancer cell division; hence, its loss of function would contribute to tumor aggression because of the loss in metabolic checkpoints(31, 32). AMPK regulates the tumor-suppressive function of the serine/threonine kinase LKB1. Ablation of LKB1 results in enhanced risk of developing gastrointestinal, lung and skin squamous cell cancers(33, 34). Moreover, AMPK is shown to inhibit PI3K/AKT/mTOR signaling, which is activated in many cancers(29, 35). Also, metabolic inhibitors such as metformin, which indirectly activates AMPK could suppress tumor growth via autophagy induction and mTOR inhibition(29, 36). Metformin is shown to inhibit the proliferation of estrogen receptor α (ERα) negative and positive breast cancer cell lines through AMPK stimulation(37). However, when tested in mice models, metformin contributes to enhanced tumor progression and increased angiogenesis, providing us with a glimpse of potential pro-neoplastic effects of AMPK activation(37).

In our study, we observed that high levels of AMPK pathway activity were associated with better outcomes in glioma, breast cancer and sarcoma (Figure 2); corroborating previous results on the tumor-suppressive function of AMPK. Conversely, the opposite is true in stomach cancer, where AMPK activation contributes to adverse outcomes (Figure 2). It has now been increasingly clear that AMPK activation can also be pro-tumorigenic(12). Double knockout of AMPKα1 and AMPKα2 in mouse embryonic fibroblasts result in impaired tumor formation(38). AMPK knockdown in pancreatic cancer cells impairs anchorage-dependent growth and reduces cell viability under glucose deprived conditions(39). AMPK signaling induces cell migration in prostate cancer cells(40) while AMPK knockdown inhibits cell proliferation and promotes apoptosis(41). In liver cancer cells retrieved from primary mouse tumors, AMPK activity is required for Myc-driven carcinogenesis(14). Taken together, these studies suggest that AMPK activation due to metabolic stress within the tumor microenvironment is crucial for the survival of cancerous cells.

Although 19 of the 21 cancers had at least one gain-of-function or loss-of-function gene that correlated with survival outcomes, glioma tumors were most influenced (Figure 2A). The consequence of dysregulated AMPK signaling was further explored in glioma, where the 24-AMPK-gene set and each of the five TFs (identified as regulators of AMPK-associated DEGs) were considered jointly for patient stratification. We observed oncogenic roles of E2F4, EZH2, FOXMI and SUZ12 – patients with high expression of these TFs had higher mortality rates (Figure 6B). Since the 24-AMPK-gene set was a positive prognostic factor in glioma where high expression of the genes was associated with better outcomes (Figure 2C), glioma patients harboring low AMPK scores and high oncogenic TF scores performed the worst. Our results are confirmed by other reports on the crosstalk between AMPK signaling and E2F4, EZH2, FOXMI or SUZ12 and their effects on oncogenic progression(42–45). Our analyses on SMAD4 in glioma revealed a likely tumor-suppressive role of the gene (Figure 6B), which is corroborated by another study demonstrating reduced SMAD4 expression during glioma tumor progression(46). When merged with the anti-neoplastic effects of AMPK activation, 5-year survival rates were improved by almost 30% compared to individuals within the low-low category (Figure 6B). SMAD4 protein expression is lost in gastric cancer cells and loss of expression in primary gastric adenocarcinomas are associated with poor survival(47). SMAD4 is also commonly inactivated in gastrointestinal cancers(48, 49). Restoration of SMAD4 expression in pancreatic cancer cells inhibits tumor function in vivo by influencing angiogenesis through decreased VEGF expression(50).

Our study has demonstrated that there is far from a single unifying role of AMPK signaling in cancer progression. Harnessing multiplatform datasets, this study provides a comprehensive depiction of how AMPK signaling is manifested in a variety of cancers. We anticipate that this repertoire of organized data would be explored by the research community to devise additional research plans aiming to better understand the roles of AMPK in cancer development. We demonstrated that the pro- or anti-neoplastic effects of AMPK activation is cancer-type dependent. Targeting AMPK for treating metabolic diseases such as diabetes has been well established. Also, the potential for targeting AMPK in cancer therapy has been elegantly reviewed(8). However, since AMPK activation is a double-edged sword, careful considerations need to be in place before AMPK can be viably deployed in clinical settings. Our study provides a comprehensive catalog of clinically actionable genetic variations which could be used for patient stratification in prospective clinical trials testing the capabilities of AMPK antagonists or agonists as potential treatments for cancer.

